# Microfluidic T-Chip enables one-step clinical-scale T-cell purification from blood products for CAR T-cell manufacturing

**DOI:** 10.64898/2026.07.30.741793

**Authors:** Maheen Rana, Sophia E. Nigrovic, Adriana Payan-Medina, Sarthak Saha, Victor R. Putaturo, Quinn E Cunneely, Remy Bell, Ezgi Antmen, Marcela V. Maus, Mehmet Toner, Magdi Elsallab, Avanish Mishra

**Author notes:** These authors contributed equally to this work (Maheen Rana, Sophia E. Nigrovic, Adriana Payan-Medina). These authors jointly supervised this work (Magdi Elsallab, Avanish Mishra). Corresponding authors: Address correspondence to Magdi Elsallab and Avanish Mishra.

## Abstract

Treatment with chimeric antigen receptor (CAR) T cells has emerged as a promising immune therapy for relapsed and refractory hematologic malignancies. The CAR T cells are manufactured in a series of steps that involve isolating T cells from the patient’s leukapheresis product, genetically modifying them to express the CAR against the target antigen, and reinfusing them into the patient. Efficient T-cell enrichment from leukapheresis products is critical to the success of these therapies. Current methods for T-cell sorting on a clinical scale involve several washing steps to remove red blood cells and platelets, followed by T-cell selection and activation. These multi-step processes result in cell loss during processing and involve several handling steps. Here, we utilize fluidically assembled micromagnetic lenses to develop a high-throughput, continuous-flow microfluidic T-cell sorter, designated as the T-Chip, for sorting magnetic bead-labeled CD3^+^ T cells in a single step. Our approach allows direct sorting of T cells in expansion media from leukopaks without any washing steps, effectively removing 99.999% of RBCs and platelets from the leukapheresis product. A single 1-inch x 3-inch T-Chip can process leukapheresis product at a throughput of 60 mL/hr and 2.56 ± 0.12 billion cells/hr. Using this optimized workflow, we demonstrate clinical-scale enrichment of highly pure CD3^+^ T cells (97.7 ± 1.3%) with high viability (97.0 ± 1.1%) and recovery (87.3 ± 14.8%) in a functionally closed manner. Downstream processing of T cells isolated using the T-Chip yielded potent anti-mesothelin CAR T cells with demonstrated anti-tumor efficacy. Overall, by exploiting precisely engineered magnetic forces and laminar flow, the microfluidic T-Chip overcomes bottlenecks caused by low throughput and enables single-step large-scale T-cell purification for the rapid development of CAR T cells.

## Introduction

Chimeric antigen receptor (CAR) T-cell therapy represents a transformative approach for cancer treatment. Guided by the extracellular domain, the chimeric receptor redirects the specificity of immune cells toward cancer cells^1^. CAR T cells targeting tumor-associated antigens such as CD19 and B-cell maturation antigen (BCMA) showed remarkable efficacy in B-cell lymphoma^2^, follicular lymphoma^3^, B-cell precursor acute lymphoblastic leukemia^4^, and multiple myeloma^5^, with overall response rates ranging from 60% to 80% in heavily pretreated patients^6^. More than a thousand active research trials are ongoing with a focus on improving CAR T-cell therapy and extending its utility to solid tumors^7^.

Currently approved CAR T-cell therapies rely on autologous cells as the starting material. The standard procedure for CAR T-cell generation involves leukapheresis to collect the patient’s peripheral blood mononuclear cells (PBMCs), enrichment and stimulation of T cells, viral transduction to introduce the CAR, and subsequent expansion^8–11^. Leukopaks contain a diverse mix of white blood cells, primarily billions of mononuclear cells, including T cells, B cells, NK cells, and monocytes, at concentrations that are over an order of magnitude higher (∼50 million cells/mL) than in peripheral blood (∼5 million cells/mL). Although leukapheresis aims to return the remaining blood constituents to the patient, residual RBCs remain at high concentrations (0.5 billion/mL), resulting in a complex blood product. Despite rapid advances in CAR T-cell therapy, the existing manufacturing process is logistically complex, involves multiple touchpoints, introduces contamination risks, and is operator-dependent. Such complexity creates a bottleneck in scaling up these therapies and making them widely accessible.

In this paper, we present a high-throughput microfluidic chip technology for clinical-scale T-cell isolation. The device enables magnetic separation of T cells in a continuous flow under an isomagnetic field, with cells only spending a few seconds in the chip. This approach represents a departure from the current magnetic cell sorters used in CAR T manufacturing, which typically operate in a batch mode and hold labeled cells against a tube or bag wall. Using this platform, we demonstrate single-step isolation of T cells from leukapheresis products without the need for pre-washing, RBC depletion, or platelet removal. We benchmark the T-Chip against column-based magnetic-activated cell sorting (MACS), a widely used isolation approach, and demonstrate superior performance with our no-wash workflow. Finally, we show the feasibility of downstream manufacturing by generating functional mesothelin-targeting CAR T cells isolated with the T-Chip.

## Results

### Microfluidic design of the T-Chip

T-Chip was constructed using lithographically defined on-chip micromagnetic lenses to efficiently deflect magnetic bead-labeled T cells from blood products at a high throughput (60 mL/hr). **Figure 1** depicts the architecture and operation of the T-Chip. The chip was designed to contain two parallel sorting channels, with the leukopak sample introduced via a co-flow of T-cell expansion media at a 1:3 ratio (leukopak:buffer) (**Figure 1A**). For sorting, T cells labeled with 4.5 µm anti-CD3 or anti-CD3/CD28 superparamagnetic Dynabeads were flowed through the chip^10^. Our results demonstrated that a single sorting channel can support a sample processing throughput of 30 mL/hr, while the co-flow expansion media is provided at 90 mL/hr. Furthermore, a 1-inch x 3-inch T-Chip supported two parallel sorting channels, achieving a total leukopak sample throughput of 60 mL/hr (**Figure 1B-C**).

**Figure 1:**
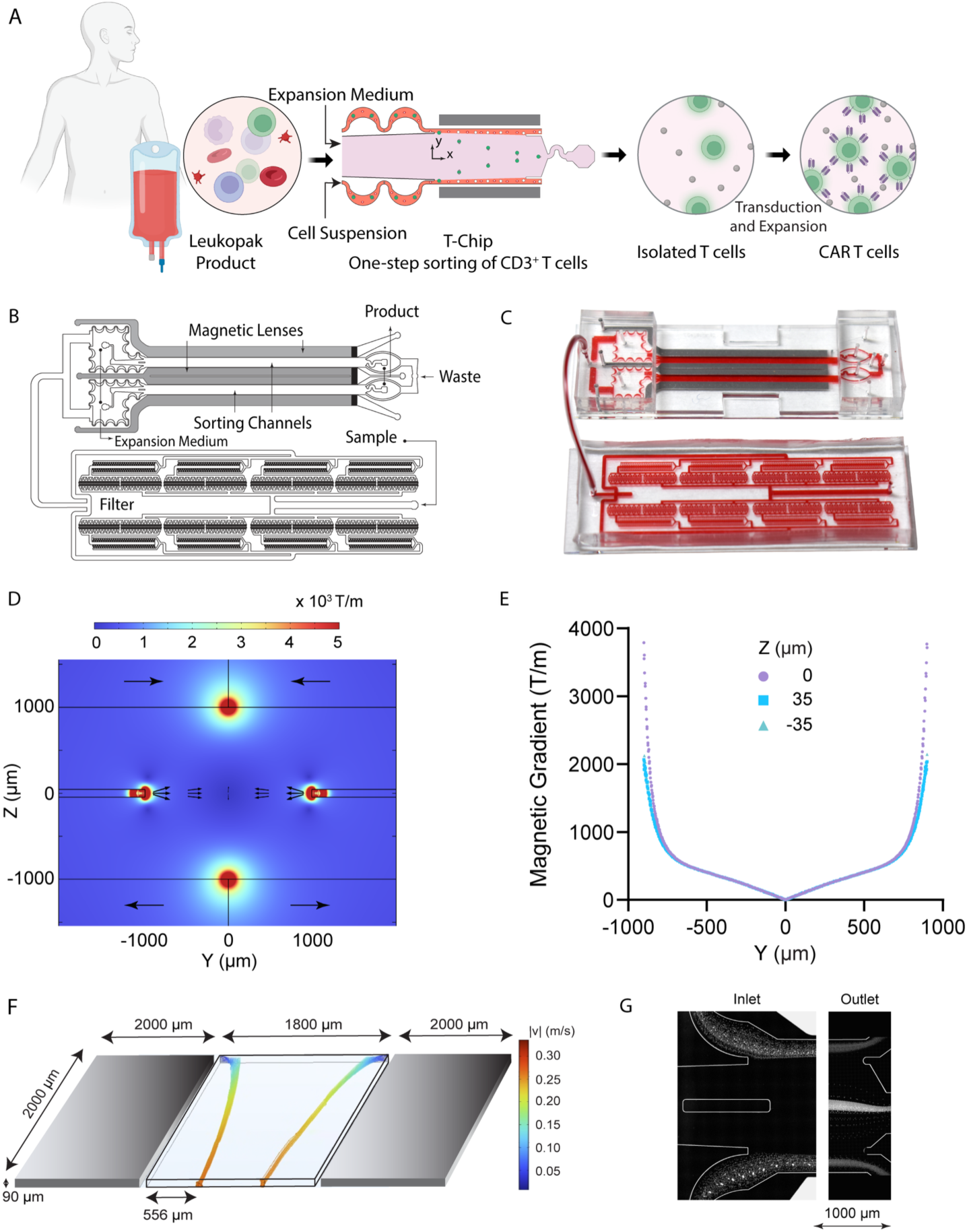
T-Chip for single-step high-throughput microfluidic magnetic isolation of T cells. (**A**) Schematic representation of the T-Chip (Created with BioRender.com). A central microfluidic sorting channel is flanked on either side by fluidically assembled magnetic lens channels, each tightly packed with high-permeability magnetic iron particles. These magnetic lenses locally intensify the magnetic field gradient to ∼3800 T/m, enabling efficient lateral deflection of labeled CD3⁺ T cells at high throughput (total cell + expansion medium flow rate = 240 mL/hr). (**B**) Layout of the T-Chip showing an upstream microfluidic pre-filtration module connected to the microfluidic sorting channels. Samples from the leukopak flow through the filtration module prior to entering the sorting channel. T-cell expansion medium is introduced through a dedicated inlet (labeled as Expansion medium), providing the main flow path for the collection of isolated T cells. (**C**) Image of the microfabricated T-Chip. (**D**) A contour plot of the magnetic gradient estimated using finite-element modeling, showing amplification by magnetic lenses. (**E**) Isomagnetic gradient across the sorting channel. The field gradient peaks near the walls of the sorting channel, coinciding with the sample stream, and decays toward the channel center. (**F**) Computational results showing deflection of an 8 µm T-cell labeled with one bead over a channel length of 20 mm, where │v│ represents particle velocity magnitude. (**G**) High-speed streak image captured in the sorting channel at the T-Chip inlet and outlet, demonstrating the targeted deflection of bead-labeled CD3⁺ T cells into the central product stream, leaving the rest of the blood cells to flow undisturbed to the waste streams.

### Fluidically assembled magnetic lenses generate an isomagnetic high-gradient force field for sorting T-cells

In our previous work, we introduced the concept of magnetic lenses to perturb magnetic field lines from permanent magnets, generating a high-gradient field in a microfluidic channel^12^. The T-Chip applies this concept to enable rapid sorting of 4.5 µm magnetic bead-labeled T cells from leukopaks (**Figure 1B**). The sorting channels were symmetrically surrounded by two parallel channels, which were densely filled with 40-µm soft-magnetic iron particles. These iron-packed channels were created by flowing a suspension of 40-µm iron particles of high magnetic permeability in 50% ethanol through the magnetic lens channel, where a filter holds the particles while allowing the suspending fluid to flow through (**Figure 1B-C**), compactly packing the channel. The supplementary material provides a detailed experimental procedure for iron filling, and **Supplementary Movie 1 explains the process**^12^. We demonstrated that the magnetic permeability contrast between the soft iron-filled lens regions and the surrounding microfluidic channel focused the magnetic field from the bulk magnets, creating steep magnetic field gradients across the sorting channel (**Figure 1D**). Permanent neodymium-iron-boron magnets were arranged in a quadrupole configuration around the sorting channel, providing a stable external magnetic field. Without magnetic lenses, the maximum magnetic field gradient in the sorting channel was 440 T/m, whereas with the magnetic lenses it increased to ∼3800 T/m (**Figure 1E**), yielding an 8.6-fold increase in the magnetic force.

In the T-Chip, the sorting channels were designed to be 1800 µm wide, 90 µm high, and 40 mm long, with parallel magnetic lens channels spaced 70 µm apart. The channel width and length were maximized to ensure ample residence time for bead-labeled T-cell deflection under the magnetic field (**Figure 1F**). Channels wider than 1800 µm were prone to collapse, while the length was maximized to 40 mm within the constraint that the T-Chip fit on a standard microscope glass slide (1-inch x 3-inch). The sorting channel height was limited to 90 µm because the magnetic field gradient became substantially nonuniform across the channel height above this threshold. We conducted three-dimensional multiphysics computational fluid dynamics (CFD) and magnetic field simulations to track deflection of bead-labeled cells (**Figure 1F**). Computational results demonstrated that at a 60 mL/hr sample throughput, cells labeled with a single magnetic bead were deflected by more than 720.3 ± 1.4 µm from the channel side wall (**Figure 1F, Supplementary Movie 2**). High-speed camera images showed that cells that are tagged with a single bead are deflected up to 745 µm from the side wall of the main channel, which is consistent with computational results (**Supplementary Figure S1**).

A pre-filter module with a 40-µm aperture was added upstream of the magnetic sorting chamber (**Figure 1B-C**). After filtration, an asymmetric serpentine channel was included to partially align the nucleated cells within the sample stream through inertial focusing (**Figure 1A, G**). This ensured uniform cell positioning in a band 60 to 220 µm from the channel sidewall before they entered the magnetic deflection zone of the sorting channel. The T-cell culture medium was introduced via a buffer inlet channel. The CD3^+^ T cells labeled with magnetic beads experienced lateral magnetic forces within the sorting channel, which guided them to the product outlet at the center of the channel, while unlabeled cells, including B cells, NK cells, monocytes, and erythrocytes, remained unaffected and flowed into the lateral waste channels (**Figure 1G**). This design enabled the direct recovery of sorted T cells into a medium suitable for downstream activation, transduction, and expansion. By accurately deflecting magnetically labeled T cells into the force-free channel core, the T-Chip enabled continuous sorting into expansion media without cell retention or device clogging (**Supplementary Movie 3)**.

### Single-step isolation of CD3⁺ T cells directly from leukopak-mimic samples using the T-Chip

To reproduce clinically relevant leukapheresis compositions, we created leukopak-mimic samples with 120–200 million mononuclear cells (MNCs) from overnight-stored blood samples using density gradient separation (*n* = 7). The hematocrit value was set to 2-3 %, and MNC cell concentration was kept at 50 million cells/mL. The mimic samples were incubated with 4.5 µm anti-CD3-conjugated beads before they were introduced directly into the T-Chip (**Figure 2A**).

**Figure 2.**
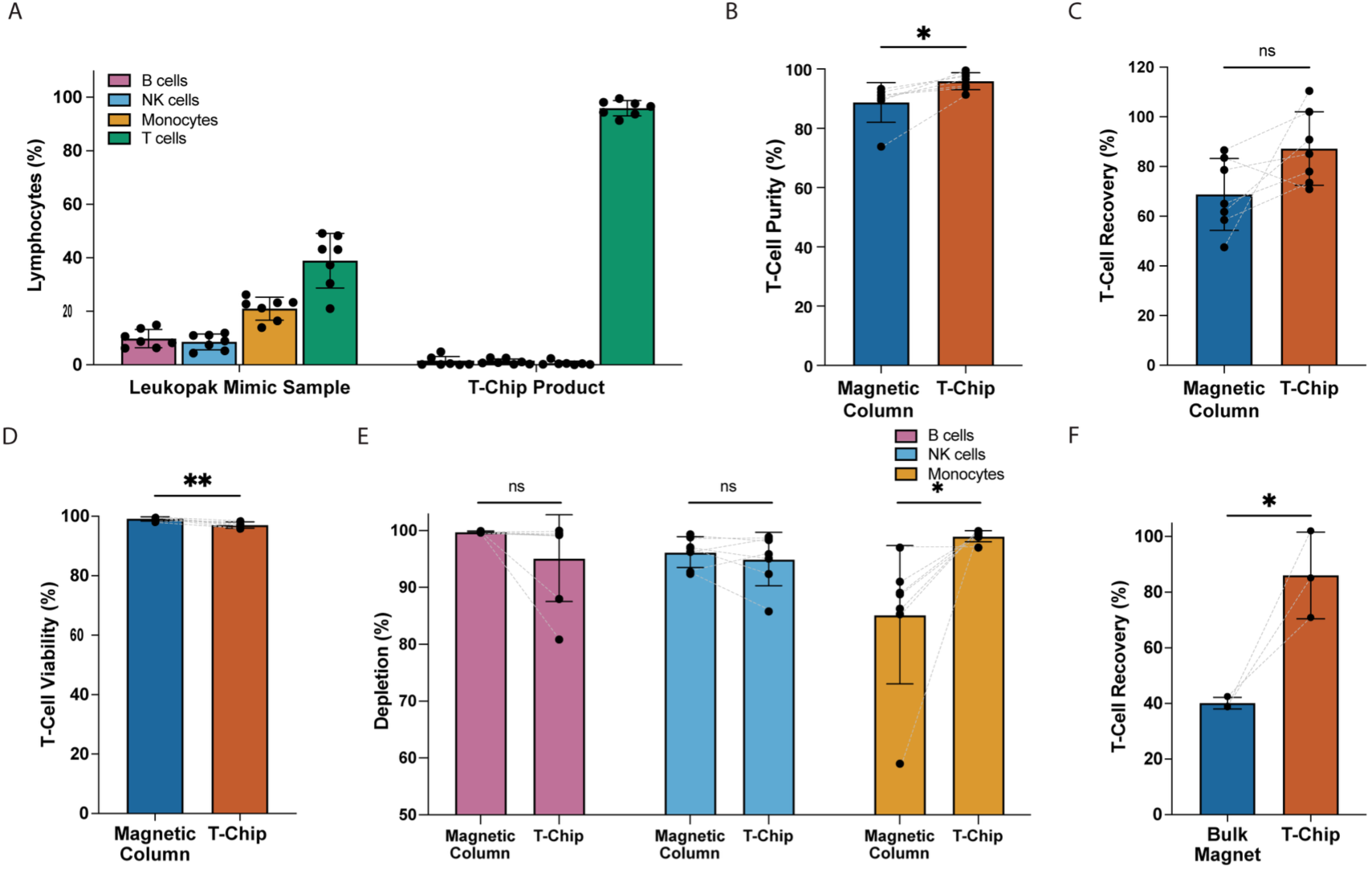
T-Chip enrichment of T cells from leukopak mimic samples. (**A**) Leukocyte distributions in the leukopak mimic sample and the T-Chip product show enrichment of T cells, with minimal contamination by B cells, NK cells, and monocytes (*n* = 7). Comparison of T-cell (**B**) purity, (**C**) recovery, (**D**) viability, and (**E**) depletion of contaminating B cells, NK cells, and monocytes between magnetic column and T-Chip (*n* = 7). **(F)** Comparison of T-cell recovery between the bulk magnet-based system and the T-Chip, showing loss of a significant number of T cells in the bulk magnet system. Data compared by paired t-test. ns = *p* > 0.05, * = *p* < 0.05, ** = *p* < 0.01. Error bars indicate standard deviation.

In seven independent experiments, the cell product obtained after T-Chip sorting showed enrichment for CD3⁺ T cells, with negligible contaminating populations (**Supplementary Figure S2A**), confirming that the deflection of bead-labeled T cells under the high-gradient magnetic field did not result in carryover of contaminating cells. In addition, the T-Chip achieved red blood cell depletion of 5.0 ± 0.3 Log₁₀, with no residual RBCs noted in hemocytometer counts in 4 out of 7 replicates. At the same time, the T-Chip achieved substantial depletion of B cells (2.0 ± 1.0 Log₁₀), NK cells (1.5 ± 0.4 Log₁₀), and monocytes (2.3 ± 0.5 Log₁₀) (**Supplementary Figure S2B-E**).

We performed a pairwise comparison of T-Chip with the current state-of-the-art Miltenyi magnetic columns for T-cell isolation. Flow cytometry of T-Chip isolated T cells demonstrated a purity of 95.9 ± 2.9%, exceeding the product purity achieved by the magnetic columns (88.7 ± 6.7%, *p* = 0.0106, **Figure 2B**). The T-Chip also yielded a T-cell recovery of 87.3 ± 14.8%, which exceeded the recovery obtained with magnetic columns (68.7 ± 14.5%, *p* = 0.0806, **Figure 2C**). The viability of T cells isolated with T-Chip was 97.0 ± 1.1%, comparable to that achieved with columns (99.1 ± 0.7%, *p* = 0.0034, **Figure 2D**). Comparison of the leukopak input and the T-Chip product indicates no significant impact on viability during processing (98.1 ± 2.4% viability for the leukopak input sample, *p* = 0.382) (**Supplementary Figure S3**). **Figure 2E** shows a comparison of the depletion efficiency of cellular impurities (B cells, NK cells, and monocytes) between the magnetic column and the T-Chip. Notably, T-Chip achieved significantly higher monocyte depletion (99.0 ± 1.0%) than the magnetic column (85.2 ± 12.0%, *p* = 0.026) (**Supplementary Figure S4**).

We also compared T-cell recovery when isolation was performed in batch magnetic racks using the DynaMag-2 magnet (*n* = 3), which is recommended in the manufacturer’s protocol for isolating cells labeled with 4.5 µm Dynabeads. In comparison to the recovery achieved by the microfluidic T-Chip, recovery was markedly reduced with bulk magnetic isolation (40.1 ± 2.1%, *p* = 0.04*, n* = 3) (**Figure 2F**), possibly due to bead detachment from cells during washing and resuspension steps. It highlights the challenges of cell isolation using bulk magnetic separation at high cell densities relevant to CAR T manufacturing. The results presented here demonstrate the T-Chip’s ability to process high-density cellular samples with excellent purity and viability exclusive of the requirement for intermediate processing steps, such as density gradient centrifugation or sequential washing.

### CAR T-cell manufacturing from the T-Chip product

Following cell isolation from leukopak mimics by the T-Chip, we evaluated the impact of the T-Chip process on downstream CAR T-cell manufacturing and function. T-Chip isolated T cells were transduced with a chimeric antigen receptor targeting mesothelin, a glycoprotein previously established as an attractive immunotherapy target due to its overexpression in several solid tumors^13–15^. Then, T cells were expanded across a standard 14-day production window (**Figure 3A**). Anti-mesothelin CAR (MesoCAR) T cells experienced an average of 112.1 ± 77.3-fold growth with 58.3 ± 9.7% surface CAR expression (**Figure 3B-D, Supplementary Figure S5**). In comparison with MesoCAR T cells produced from magnetic column-isolated T cells, T-Chip-derived MesoCAR T cells demonstrated enhanced surface CAR expression (58.3 ± 9.7% vs 50.3± 7.8%, *p* = 0.0184, **Supplementary Figure S5A**) and similar fold expansion (112.1 ± 77.3 vs 154.4 ± 72.6, *p* = 0.1484, **Supplementary Figure S5B**). Notably, cytotoxic CD8^+^ T cells were enriched in the T-Chip output (34.4 ± 10.5% vs 26.6 ± 7.5%, *p* = 0.0116, **Supplementary Figure S6**), in MesoCAR T cells (50.7 ± 6.7% vs 34.8 ± 4.8%, *p* = 0.0020, **Supplementary Figure S7**), and in the 13-day expansion of T-Chip product (65.5 ± 23.8 % vs 47.7 ± 17.2% *p* = 0.0024, **Supplementary Figure S8**).

**Figure 3.**
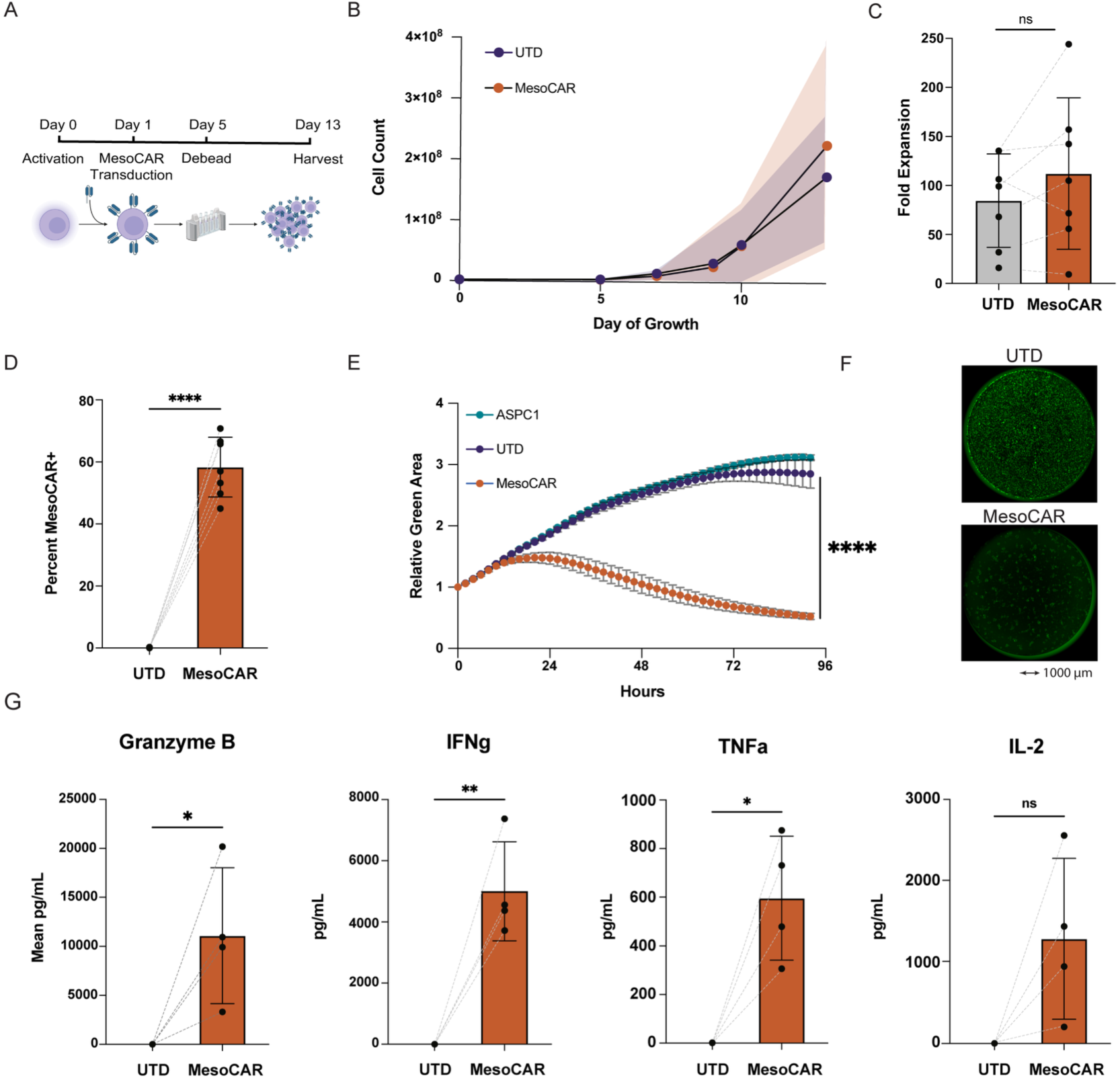
T-Chip-selected T cells produced functional anti-mesothelin CAR T cells (MesoCAR). **(A)** Schematic of MesoCAR T-cell production timeline. **(B)** Growth (*n* = 7), **(C)** fold expansion (*n* = 7), and **(D)** surface MesoCAR expression of untransduced (UTD) vs MesoCAR T cells at the end of production (*n* = 7). **(E)** ASPC1-GFP tumor killing by UTD vs MesoCAR T cells (*n* = 5) in technical triplicates as assessed by IncuCyte assay and compared by two-way ANOVA. **(F)** Activated MesoCAR T-cell clustering around ASPC1-GFP tumor cells in the IncuCyte compared to UTD. **(G)** Granzyme B, IFN γ, IL-2, and TNFα cytotoxic cytokines measured via Ella Automated ELISA from the supernatant following 4 days of T-cell co-culture with ASPC1-GFP tumor cells (*n* = 4). Conditions compared by paired t-test. ns = p > 0.05, * = p < 0.05, ** = p < 0.01, **** = p < 0.0001.

Anti-tumor efficacy was likewise unimpaired compared to CAR T cells from column-isolated T cells. When co-cultured with the mesothelin-expressing pancreatic adenocarcinoma cell line (ASPC1), T-Chip-derived MesoCAR T cells successfully eliminated the tumor within 96 hours (**Figure 3E, 3F**). MesoCAR T-cell functionality was further confirmed by evaluating their secretory function in a subset of four independent experiments (*n* = 4). Following co-culture with ASPC1 tumor cells, elevated levels of effector cytokines granzyme B (11090 ± 6937 pg/mL vs 0.9 ± 1.4 pg/mL UTD, *p* = 0.0494), IFN gamma (5004 ± 1618 pg/mL vs 0.3 ± 0.3 pg/mL UTD, *p* = 0.0085), and TNF alpha (596 ± 255 pg/mL vs 0.5 ± 0.5 pg/mL UTD, *p*=0.0185) were present in the co-culture supernatant (**Figure 3G**). Higher concentrations of activation cytokine IL-2 were also observed in MesoCAR T-cell co-cultures, with some donor variability (1288 ± 988 pg/mL vs 2.7 ± 0.9 pg/mL UTD, *p* = 0.0801, **Figure 3G**). In summary, T cells isolated using the T-Chip recapitulated expected MesoCAR T-cell manufacturing dynamics and effector function.

### Functionally closed clinical-scale T-cell isolation from large-volume leukapheresis products

Based on our small-scale experiments, we demonstrated T-cell isolation and CAR T-cell manufacturing from leukopak-mimic samples containing 120-200 million MNCs. To more closely recapitulate the clinical manufacturing workflow, in which approximately 1 to 2 billion MNCs are typically processed to ensure recovery of more than 100 million T cells as starting material^16^, we next evaluated T-Chip performance at a 10-fold larger scale. In these experiments, the T-Chip successfully processed high-cellularity samples containing an average of 1.71 ± 0.08 billion nucleated cells (*n* = 3). Cells collected via leukapheresis were processed in a single T-Chip in 40 minutes at 2.56 ± 0.12 billion cells/hr without clogging, ensuring pure T-cell isolation with high recovery (**Figure 4**). The nucleated cell density in these samples ranged from 28 – 62 million cells/mL (48.67 ± 18.24 million cells/mL) (**Figure 4A**). The leukopaks were processed directly from the collection bag in a functionally closed manner to demonstrate clinical translatability, as shown in **Figure 4B** and in **Supplementary Figure S9**. As part of this closed circuit, beads were added to the leukopak bag through a sterile connector, incubated for 60 minutes, and cells were then pumped with a peristaltic pump from the leukopak bag to the T-Chip via a fluidic dampener, which was pressurized to 8 PSI by a digital pressure regulator. This regulator was also used to pressurize the bottle containing the T-cell media buffer. An air filter was applied to keep the air passing through it sterile. The flow dampener minimized the peristaltic pulsations and enabled precise fluid transfer for T-cell enrichment. The presented closed system allowed for cell isolation without contact with air or manual handling.

**Figure 4.**
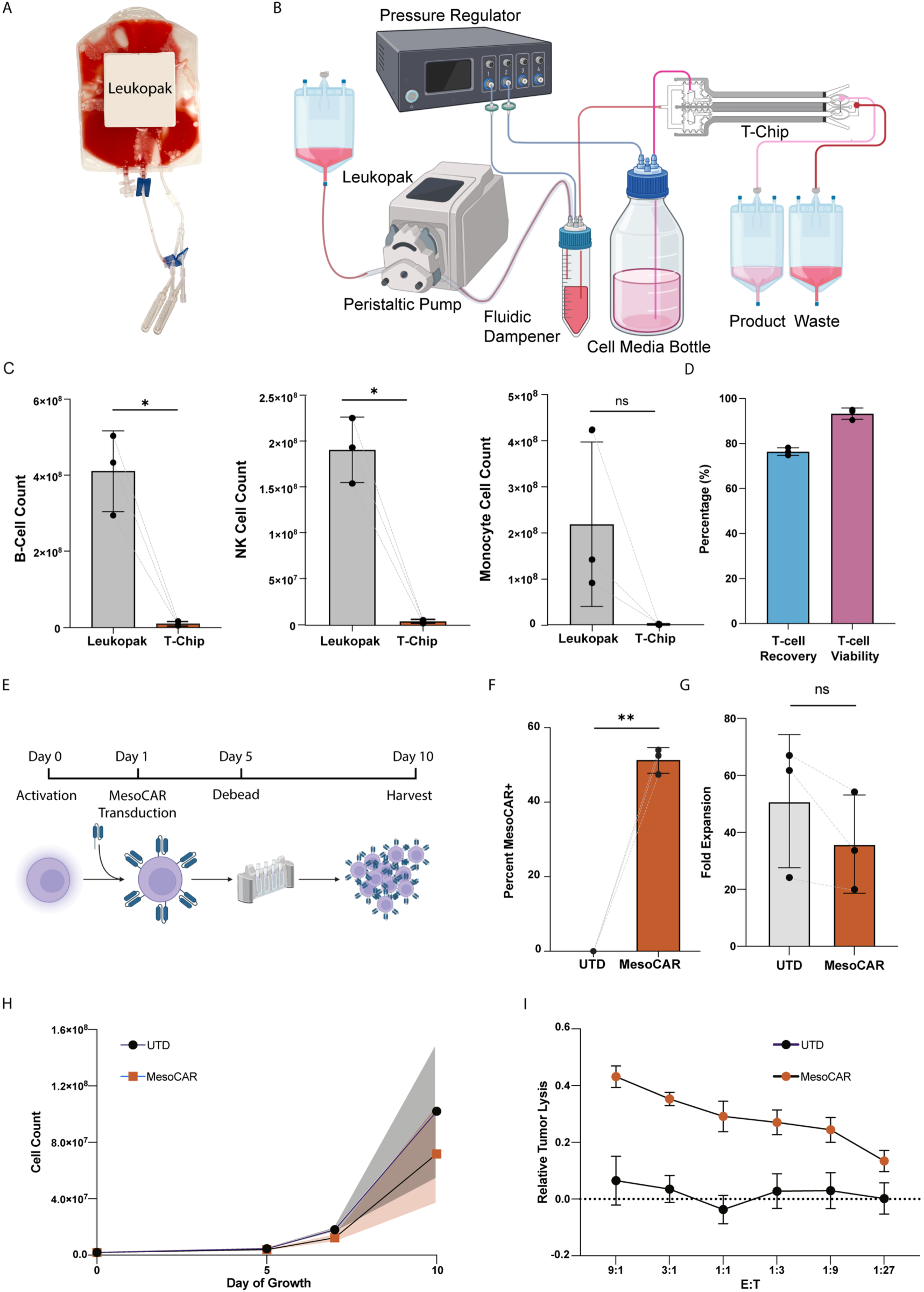
T-cell isolation from clinical-scale leukapheresis samples. (**A**) Photograph of a representative leukopak sample prior to processing. This image was edited to cover the donor-specific label. (**B**) Schematics of a closed-loop circuit for leukopak processing using the T-Chip, including a fluidic dampener and digital pressure control. Beads are added to the leukopak bag through a sterile connector and incubated for 60 minutes to keep the process closed. Cells are pumped from the leukopak bag into the T-Chip with a peristaltic pump, and the flow rate is kept steady by a fluidic dampener controlled by a digital pressure regulator. The same regulator also pressurizes the bottle with T-cell media buffer. This configuration minimized pulsation, enabled controlled T-cell enrichment, and eliminated air exposure or manual intermediate handling (Created with BioRender.com). (**C**) Depletion of B cells, NK cells, and monocytes from leukapheresis by T-Chip (*n* = 3). (**D**) Recovery and viability of T cells isolated using T-Chip from leukopak samples (*n* = 3). (**E**) Clinically relevant production timeline for MesoCAR T cells manufactured from T cells isolated by one-step sorting from a leukopak. (**F**) Surface expression of the MesoCAR on untransduced (UTD) and MesoCAR T cells after 10 days of production (*n* = 3). (**G-H**) Fold expansion (**G**) and growth (**H**) of MesoCAR T cells post T-Chip isolation (*n* = 3). (**I**) Relative lysis of luciferase-expressing ASPC1-GFP cells by MesoCAR T cells and donor-matched UTDs co-cultured in technical triplicates for 24 hours at varying effector (E): target (T) ratios (*n* = 3). Column data compared by paired t-test, with ns = p > 0.05, * = p < 0.05, ** = p < 0.01.

The leukopak composition consisted of 57.1 ± 6.0% CD3⁺ T cells, which were enriched to 97.7 ± 1.3% (*n*=3 independent experiments) (**Supplementary Figure S10**). **Figure 4C** shows efficient depletion of contaminating cells in the leukopak using T-Chip, achieving 1.7 ± 0.2 Log_10_ B-cell depletion, 1.6 ± 0.2 Log_10_ NK cell depletion, and 2.2 ± 0.6 Log_10_ monocyte depletion, underscoring efficient microfluidic enrichment of T cells and removal of contaminating immune subsets in the product. Despite processing such a large quantity of cells, which were previously stored for 24 hours at room temperature, the recovery was 76.4 ± 1.7%, and viability was 93.3 ± 2.5% (**Figure 4D**).

MesoCAR T cells were manufactured from T cells isolated by T-Chip, employing a production timeline that mimicked the clinical, large-scale generation of CAR T cells (**Figure 4E**). Upon MesoCAR T-cell harvest on day 10, MesoCAR T cells had expanded, on average, 35.9 ± 17.2-fold, with 51.2 ± 3.5% of cells in culture expressing the anti-mesothelin CAR (**Figure 4F-H**). The large-scale MesoCAR T cells likewise successfully eliminated mesothelin-expressing ASPC1-GFP tumor cells within a 96-hour co-culture assay (**Supplementary Figure S11**). To further test the MesoCAR T cell potency, MesoCAR T cells were co-cultured at varying effector (CAR):target (ASPC1) cell ratios and killing of the luciferase-expressing ASPC1 tumor cells was assessed after 24 hours. As expected, the MesoCAR T cells demonstrated mesothelin-targeted cytotoxicity at all E:T ratios tested (p < 0.0001, **Figure 4I**). Together, these findings demonstrate that a single T-Chip can process clinically relevant, high-cellularity, 24-hour-preserved leukopak samples in a functionally closed format without clogging, while maintaining high T-cell purity, recovery, and viability. Importantly, isolated cells supported efficient generation of functional CAR T cells with potent, target-specific anti-tumor activity.

## Discussion

Cell isolation from complex blood products is a critical step for robust cell therapy production^12^. A typical CAR T-cell production begins with 50 to 100 million T cells, selected from 1 to 2 billion input MNCs^8^. Various microfluidic technologies have been studied to obtain T-cell enrichment. Of note is acoustophoresis^17–19^, but it mainly sorts cells based on size and deformability^20^. Inertial microfluidics^21–27^ exploiting hydrodynamic lift and drag forces has also been used to isolate target cells based on size and deformability^26,28,29^. However, these tools have limited utility for sorting T cells due to their overlap in size and deformability with other lymphocytes. Given that blood cell populations substantially overlap in size and morphology, cell-surface markers provide the most practical basis for T-cell enrichment^30^. Accordingly, current methods for cell sorting include fluorescence-activated cell sorting (FACS) or magnetically activated cell sorting (MACS). Although FACS enables multiparameter cell sorting, GMP-compliant FACS systems are limited in throughput and are therefore mostly used as a secondary purification step after initial MACS enrichment^31^. Finally, magnetophoretic approaches remain the clinically approved gold standard for CAR T manufacturing because antibody-conjugated magnetic beads can be used to obtain marker-specific T-cell populations, ensuring high purity, while operating at a higher throughput than FACS^8,32^.

MACS technologies utilize superparamagnetic beads conjugated with T-cell-specific antibodies (anti-CD3, anti-CD4, or anti-CD8) to achieve positive selection of T cells^33–38^. Alternatively, negative selection can be used to deplete non-T-cell populations by using CD16/CD56/CD19/CD14-labeled beads to remove NK cells, B cells, and monocytes^39^. Still, conventional MACS systems, such as magnetic columns, bulk magnets, and the more advanced and automated CliniMACS Plus systems, necessitate multiple washing steps for upstream product cleanup to remove plasma, RBCs, and platelets, followed by multiple intermediate batch processing steps during magnetic T-cell enrichment, leading to prolonged processing times and system-wide variability^10,11,40^. Using centrifugation-based washing steps at low or no braking increases the processing time for large-scale leukapheresis samples to 4-6 hours while yielding recoveries of only ∼50%^41,42^. Furthermore, the presence of non-T-cell impurities, such as monocytes, leads to T-cell apoptosis and has been shown to interfere with magnetic bead binding to T cells, thereby ultimately impairing T-cell activation^39^. The presence of monocytes also correlates with lower T-cell expansion and transduction efficiency^34,43–45^. In anti-CD19 CAR T-cell manufacturing, contaminating CD19^+^ leukemic cells may also be transduced, mediating antigen masking and immune evasion^46^.

Microfluidic MACS technologies, including the ^LP^CTC-iChip developed in our lab, enable high-throughput cell isolation with minimal loss^12,47^. The ^LP^CTC-iChip can process over 6 billion cells per hour, depleting more than 99.99% of contaminating cells, thereby creating an ultrapure population of rare tumor cells using negative selection^17–19^. However, this system still requires microfluidic washing steps across separate devices to remove RBCs, platelets, and plasma. Here, we presented a standard glass slide-sized positive-selection magnetic chip that uses fluidically assembled magnetic lenses to sort CD3^+^ T cells directly from clinically relevant leukapheresis material at a high flow rate of 60 mL/hr, while removing unwanted WBCs and RBCs in a single step. Using this approach, we demonstrated that the T-Chip successfully selected CD3^+^ T cells from clinical-scale leukapheresis samples to an average purity of 97.7 ± 1.3% without clogging and with minimum processing requirements, limited to mixing of cells with the beads and incubation.

Despite processing 24-hour preserved samples, after T-Chip processing, cells maintained high viability (93.3 ± 2.5%) and consistently lower contamination was noted as compared to the gold standard method using magnetic columns^10,11,48–50^. Furthermore, downstream manufacturing of anti-mesothelin CAR T cells confirmed T-cell functionality, showing no deleterious impact of the T-Chip on proliferation, transduction, antigen recognition, and target-specific killing, as evidenced by multiple functional killing assays and cytokine production capabilities.

T-Chip relies on photolithography to create channels for magnetic lenses at precisely defined distances from the sorting channel. That way, magnetic lenses can be integrated directly into the microfluidic device, generating a magnetic field right next to the sorting channel without obstructing fluid flow. This approach provides flexibility in device and channel architecture by separating magnetic field generation from microfluidic design. Bead-tagged T cells are deflected toward the channel core, where they experience a force-free region owing to the isomagnetic field gradient generated by magnetic lenses that vanishes at the center of the sorting channel. The presented arrangement of magnetic force naturally avoids T-Chip clogging and allows for processing billions of cells continuously. To achieve the isomagnetic field gradient, the channel of the magnetic lens must be the same height as the sorting channel (90 µm). While metals can be patterned using sputtering, electron beam evaporation, and thermal evaporation, the fabrication of 90 µm-thick features using these methods presents a challenge, and the resulting alignment with sorting channels further adds to the complexity of manufacturing and scalability.

Our proposed concept differs from previously described magnetic isolation strategies. Adams et al.^51^, for example, described a multiplex magnetic sorter for *Escherichia coli* that requires magnetic beads of different sizes to select strains of variable cell size. Poudineh et al.^52^ also designed a nanoparticle-based platform for magnetic ranking cytometry, which uses surface-functionalized magnetic nanoparticles to profile and characterize circulating tumor cells based on the expression of surface markers. This approach, however, was primarily designed for the phenotypic characterization of rare cells. In another study, Pamme and Wilhelm^53^ and Robert et al.^54^ presented a free-flow magnetophoresis approach that sorts up to 0.5 million cells per hour. Our group also presented a cell sorter based on a quadrupole magnetic arrangement^23,55,56^, but it also showed limited ability to process large blood volumes and elevated cell concentrations (<5 million cells/mL), as required for processing heterogeneous leukopak compositions.

Importantly, compared with magnetic columns and bulk magnets, where bead-labeled cells are retained against a porous matrix or bag wall, the T-Chip continuously deflects tagged cells toward the channel center, away from channel walls into a clean buffer. This non-contact separation mechanism also makes it less likely to get clogged when dealing with high cell concentrations, such as those found in leukapheresis products, and may have contributed to the higher yield and purity observed with the T-Chip compared to the magnetic column. The magnetic column also must undergo several wash cycles to remove unlabeled cells, necessitating a batch-processing workflow; even automated platforms such as CliniMACS Plus require washing in successive, individual steps. In contrast, the T-Chip essentially removes all red blood cells and unwanted B cells, NK cells, and monocytes in a single step and in a continuous flow, resulting in a pure product, which is critical for the later production of viable CAR T cells. In addition, conventional CAR T manufacturing processes (e.g., CliniMACS) perform T-cell selection based on specific subpopulations, such as CD4^+^ and CD8^+^, using nanobeads conjugated to the corresponding antibodies. While these approaches yield defined T-cell populations, they require an additional step to further activate the selected T-cells in a separate process using either soluble or bead-conjugated antibodies. Our T-Chip platform, on the other hand, can perform selection and activation in a single step by using anti-CD3/CD28 beads.

To validate the clinical GMP applicability, we implemented the T-Chip platform in a functionally closed configuration with large-volume leukapheresis samples. Closed platforms are critical for minimizing the risk of contamination and achieving regulatory compliance, as well as minimizing operator errors^10,11^. We demonstrated a two-stage closed-loop flow strategy in which samples from the leukapheresis product were pumped by a peristaltic pump through a pressurized fluidic dampener to minimize peristalsis-induced pulsations. This setup maintained a sterile environment, providing a stable, uninterrupted flow to enrich CD3^+^ T cells from the leukopak bag without manual handling.

One limitation of the current device architecture is the need for precise alignment of the microfluidic chips with magnets to achieve an optimal magnetic field gradient. In the future, we plan to address this constraint by developing an alignment-free magnetic circuit that would simplify device setup and reduce variability. This platform is currently designed for micrometer-scale magnetic beads and is not compatible with cells labeled with nanometer-sized particles. Finally, this work was performed using blood products from healthy donors. Future efforts will aim to validate this platform using patient-derived leukapheresis material.

In conclusion, this work establishes fluidically assembled micromagnetic lenses as a powerful tool for clinical-scale T-cell enrichment from complex blood products using micron-sized magnetic beads. T-Chip results in direct isolation of highly pure and viable T cells from complex leukapheresis products without upstream washing, intermediate handling, or cell retention. These findings demonstrate that precisely engineered magnetic forces in a microfluidic channel can transform complex biological inputs into manufacturing-ready cell products, opening new opportunities for rapid production of cell therapies.

## Experimental Section

### Integrated T-cell sorting workflow

To enable fast and scalable enrichment of T cells appropriate for downstream CAR T manufacturing, we have established a closed, microfluidic sorting platform that groups magnetic sorting with direct collection into expansion media. The sorting workflow commenced with either clinical-grade leukopaks (*n* = 3) (**Figure 4),** acquired from healthy donors by leukapheresis, or leukopak mimic samples (*n* = 7) prepared in-house by Ficoll density gradient separation of whole blood **(Figure 2)**. These mimic samples normally contained ∼120-200 million cells and are maintained at ∼2 % hematocrit. Whole blood and leukapheresis products from healthy donors were procured from Research Blood Components, LLC (Brighton, MA).

In the Leukopak samples (*n* = 3), 60-minute incubation with anti-CD3-conjugated Dynabeads was performed, while in the Leukopak mimic samples (*n* = 7), incubation was performed for 20–30 minutes, with both steps carried out at room temperature with gentle shaking. The bead-labeled cells were then loaded into the microfluidic T-Chip sorter for magnetic isolation. The sorted mL product is captured in T-cell expansion media, eliminating the need for further transfer or purification steps.

### T-Chip design, fabrication and assembly

The microfluidic T-cell sorting device features a magnetic sorting channel surrounded by magnetic lenses (filled with soft-magnetic iron) and a pre-filter module upstream of the main sorting channel. The main sorting channel is 1800 µm wide and 90 µm high, with the product deflected toward the channel center and the waste following its original path to the two side streams. The cells tagged with beads enter the pre-filtration module to remove larger particles and aggregated cellular debris, then flow at 60 mL/hr through a connected tube into the sample inlet, while a center stream of T-cell expansion medium is fed at 180 mL/hr through the buffer inlets. The T cells were deflected in the sorting channel by carefully aligning the T-Chip with a series of neodymium-iron-boron permanent magnets (class N52) arranged in a quadrupolar configuration and mounted in an aluminum holder.

Conventional PDMS soft lithography techniques were used to fabricate the T-Chip sorting devices. SU-8 50 photoresist (Kayaku Advanced Materials) was spin-coated onto a 4-inch silicon wafer at 1900 rpm for 30 seconds to produce a master mold. Using photolithography, microchannel patterns were transferred to the resist layer by exposure to ultraviolet light (365 nm) through a high-resolution chrome mask and then developed with a SU-8 developer (Baker BTS-220) to reveal the channel structures.

PDMS (Sylgard 184, Dow Corning) was prepared by mixing the base and curing agent in a 7:1 ratio. The mixture was poured onto the mold and degassed in a vacuum chamber before being cured at 65°C for 8–12 hours to form the device layer (∼1 mm thick). To hold the press-fit tubing in place, we made separate 3-mm-thick rectangular PDMS interface pads (35 mm × 15 mm) on a plain mold and attached them to the primary device after treating them with oxygen plasma. We then punched inlet and outlet holes (1.2 mm in diameter) using a sterile biopsy punch. The PDMS layers were permanently bonded to a pre-cleaned glass substrate using plasma treatment. We then performed thermal post-curing at 85°C for 10 minutes on the bonded device before finally baking it for 3 hours at 150°C to improve its mechanical stability and prevent deformation under flow.

### Magnetic beads preparation and incubation

CD3-Dynabeads (4.5 µm) (Thermo Fisher Scientific) were used to enrich and select CD3^+^ T cells. The manufacturer’s protocol was followed to wash the Dynabeads on a magnetic stand using a PBS buffer supplemented with 0.5% BSA and 2 mM EDTA. The washing steps were repeated three times. The beads were resuspended in the wash buffer to the original volume after the wash step, then added to the samples at a 1.2:1 (beads to cells) ratio. The mixture was positioned on a gentle shaker at room temperature to allow for efficient labeling.

### Column-based T-cell selection

Buffy coats were washed in MACS buffer (0.5% bovine serum albumin (BSA) and 2 mM EDTA in Dulbecco’s Phosphate Buffered Saline (DPBS)) prior to being tagged with CD4 and CD8 MicroBeads (Miltenyi Biotec) as per the manufacturer’s protocol. Cells were resuspended at up to 10^8^ cells per 500 µL buffer and loaded onto LS Columns (Miltenyi Biotec) mounted on a QuadroMACS Separator (Miltenyi Biotec). CD4^+^ and CD8^+^ cells were then collected in MACS buffer, and T-cell purity was assessed by flow cytometry.

### Post-sorting analysis

After sorting, T-cell purity, yield, and viability, as well as erythrocyte removal, were evaluated. The “input,” “product,” and “waste” fractions were collected separately and analyzed by flow cytometry to quantify T-cell purification and the removal of unwanted immune cell fractions, particularly monocytes, B cells, and NK cells. Cell viability was acquired by trypan blue staining. The number of nucleated cells was attained by manual counting using a hemocytometer (Haemocyte) with DyeCycle Green, a DNA-binding fluorescent dye that selectively stains nucleated cells. Erythrocyte depletion was counted using bright-field hemocytometry, and the erythrocyte counts of the initial and product fractions were evaluated.

### Flow cytometry

Cells were stained in chilled FACS buffer (2% fetal bovine serum (FBS, Corning) in DPBS) supplemented with 10% Brilliant Stain Buffer Plus (BD Biosciences) and 5% Human BD Fc Block (BD Biosciences). The following conjugated mouse anti-human antibodies were used for staining: CD2 APC/Fire750 (clone RPA-2.10, BioLegend, 1:100); CD3 APC-H7 (clone SK7, BD Biosciences, 1-3:100); CD3 APC/Fire750 (clone SK7, BioLegend, 1-3:100); CD4 BV421 (clone SK3, BioLegend, 1:200); CD5 FITC (clone UCHT2, BioLegend, 1:100); CD8 V500-C (clone SK1, BD Biosciences, 3:100); CD14 APC (clone MφP9, BD Biosciences, 1:200); CD19 PE (clone SJ25C1, BioLegend, 1:100); CD34 APC (clone 8G12, BD Biosciences, 1:100); CD45 PE-Cy7 (clone 2D1, BioLegend, 1:50); CD56 BV711 (clone HCD56, BioLegend, 1:50). Cell viability was determined by 7-amino-actinomycin D staining (BioLegend, 1:100) added prior to flow cytometry analysis on a BD FACSLyric Flow Cytometer (BD Biosciences). **Supplementary Figures 12**-**14** show a representative flow cytometry gating strategy for the immune cell characterization panel in the Leukopak input, the processed T-Chip product, and the MesoCAR T-cell characterization panel, respectively.

### CAR T-cell production

Selected T cells were activated with 1:100 T-cell TransACT (Miltenyi Biotec) and grown in TexMACS Medium (Miltenyi Biotec) supplemented with 5% heat-inactivated FBS, 1X Penicillin-Streptomycin (Gibco), and 50 IU/mL IL-2 (Miltenyi Biotec). Anti-mesothelin CAR was lentivirally transduced one day after activation, and the activation reagent and beads were removed on day 5. T cells were cultured in the small-scale production for a total of 13 days, while in the large-scale production, T cells were harvested on day 10 to reflect a more clinically relevant process. MesoCAR surface expression as a measure of transduction was assessed by flow cytometry at the end of production.

### T-Chip killing assays

For the longitudinal IncuCyte cytotoxicity assay, ASPC1-GFP cells in R10 medium (10% FBS, 1% Penicillin-Streptomycin (Gibco) in RPMI Medium 1640 (Gibco)) were seeded in a flat-bottomed 96-well plate at 80% of desired confluency and allowed to attach overnight prior to co-culture. MesoCAR and donor-matched untransduced (UTD) cells were also thawed the day prior to co-culture and rested overnight in R10 containing 50 IU/mL IL-2. On the day of co-culture, MesoCAR expression was assessed via flow cytometry and normalized across all donors. MesoCAR and UTD cells were then gently added to the adherent ASPC1-GFP cells at a 1:1 effector:target ratio in technical triplicates. ASPC1-GFP tumor growth, as indicated by green fluorescence area, was measured by IncuCyte Live-Cell Analysis System (Sartorius) for 4 days.

For the luciferase-based cytotoxicity assay, normalized MesoCAR and UTD cells at 10^6^ CAR+ per mL were sequentially diluted three-fold in a 96-well round-bottomed plate. ASPC1-GFP cells expressing firefly luciferase were added to each well at 10^6^ tumor cells/mL to achieve a range of effector:target ratios. T cells and tumor were co-cultured for 24 hours prior to luciferase activity detection by Bio-Glo Luciferase Assay System (Promega) and BioTek Synergy Neo2 plate reader (Agilent Technologies, Gen5 2.09 software). Fluorescence intensity inversely correlated with CAR T-cell cytotoxicity.

### Ella cytokine analysis

Cell-free supernatants from longitudinal IncuCyte killing assays were harvested at the end of the co-incubation period and clarified by brief centrifugation at 2000xg to remove residual cells and debris. Samples were diluted 1:6 in the manufacturer-provided sample diluent, and 25 µL of diluted supernatant were loaded per sample inlet onto a Simple Plex Cell Activation Panel 1 cartridge (Bio-Techne), together with the supplied wash buffer. Cytokine quantification was performed using the Ella automated microfluidic immunoassay platform (Bio-Techne) following the manufacturer’s protocol. Concentrations were calculated using factory-calibrated standard curves and reported as pg/mL.

### Statistical analysis

Statistical analysis was performed using GraphPad Prism 10. Data points and error bars represent the mean value and standard deviation, respectively. Significance values reflect the results of a paired *t*-test between matched samples, in which *p* < 0.05 was considered significant.

## Acknowledgments

This study was supported by NIH Grant K25HL169816 (A.M.).

## Author Contributions

M.R., A.M., and M.E. conceived and designed the study. M.R., A.M., and M.E. developed the methodology. M.R., S.N., and A.P.-M. performed the investigations. M.R., S.N., A.M., M.E., M.T., and M.V.M. contributed to data visualization and interpretation. S.S. performed computational simulations. A.M. and M.E. supervised the project. M.R. and A.M. wrote the original draft of the manuscript. M.R., A.M., M.E., S.N., A.P.-M., S.S., V.R.P., Q.E.C., R.B., E.A., M.V.M., and M.T. reviewed and edited the manuscript. V.R.P., R.B., and E.A. provided technical support. All authors discussed the results and approved the final version of the manuscript.

## Data Availability Statement

The research data supporting this publication are available from the corresponding author upon request.

