## Supplemental Figures S1-S14 for "Microfluidic T-Chip enables one-step clinical-scale T-cell purification from blood products for CAR T-cell manufacturing"

Microfluidics, T Cell Isolation, Immunomagnetic Separation, Cell Therapy Manufacturing, Leukapheresis, Cancer Immunotherapy

### COMSOL simulation

A 3D computational model of the channel and the magnetic lens was developed in COMSOL Multiphysics. To optimize memory and computational resources, the physics was decoupled into three separate studies. First, a stationary study was used to compute the magnetic field using the "*Magnetic Fields, No current*" interface. For the iron channel, the relative permeability was assigned as 10,000. For the neodymium magnets, the relative permeability was 1.05, and the remanent flux density was 1.42 T, while for the surrounding media, the relative permeability was 1. Second, the fluid flow in the channel was simulated using a stationary *"Laminar Flow"* physics interface. To simplify the computation, a fully developed unidirectional flow profile was assumed, and a standard no-slip boundary condition was applied to the lateral channel walls. Finally, a time-dependent study was performed using the *"Particle Tracking for Fluid Flow*" interface. Under this study, drag force (derived from *Laminar Flow*) and magnetophoretic forces (derived from *mfnc*) were applied to the particle. To simulate the payload of a magnetic bead, cells conjugated with 1 magnetic bead were assigned a relative permeability of 1.022. A particle size of 10 µm and a density of 1050 kg/m3 were used.

The model was discretized using a swept mesh with an element size of 1 µm. Due to computational memory constraints imposed by the fine mesh resolution, the simulation was divided into two sequential 20 mm domains. Particles were initially released at random coordinates within the first 200 µm of the channel wall to mimic experimental inlet conditions. To accurately capture the total cellular deflection over 40 mm, the outlet spatial coordinates of the particles from the first 20 mm simulation were extracted and applied as the initial inlet positions for the subsequent 20 mm simulation.

### Iron particle filling in T-Chip

The T-Chip magnetic lenses were assembled by flowing a suspension of 40 µm soft-magnetic iron particles and holding them against an array of microfilters (**Supplementary Movie 1**). A 50 wt% suspension of iron particles in ethanol was prepared by gentle vortexing to ensure a uniform mixture. The suspension was promptly drawn into a 1 mL syringe, ensuring that the particles were collected from the bottom of the tube to obtain a high particle concentration. The syringe was connected to an assembly consisting of a Luer lock connector, a silicone tube, and a Tygon tube (0.02 inches ID), and another tube was inserted into the chamber's outlet to flush extra ethanol. The syringe was held upright to remove any trapped air and fill the microchannels with ethanol. The syringe was then inverted to allow the iron particles to enter the chamber via the tubing assembly. Constant pressure was applied to push the particles through the microchannel until the flow front reached the opposite end of the chamber. At the end of the filling process, the tubing was clamped off, and a slight negative pressure was created by gently pulling back on the syringe before disconnecting it to prevent leakage or backflow of particles. The process was repeated for each of the remaining chambers in turn. Finally, the device was placed on a heating plate at 50 °C for 1 hour to allow the packed particle structures to solidify.

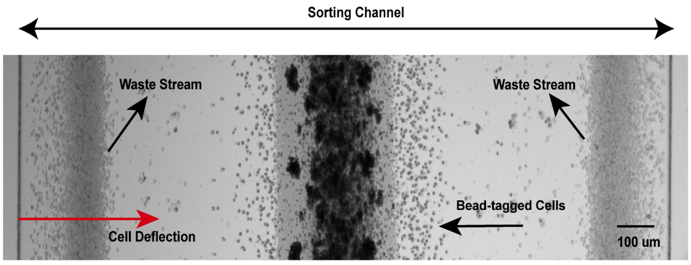


**Figure S1. Magnetic deflection of T cells labeled with beads in the T-Chip sorting channel.** A representative high-speed image of cells labeled with magnetic beads, which are magnetically deflected up to 745 µm from the sidewall of the channel and move toward the centerline of the T-Chip's sorting channel.

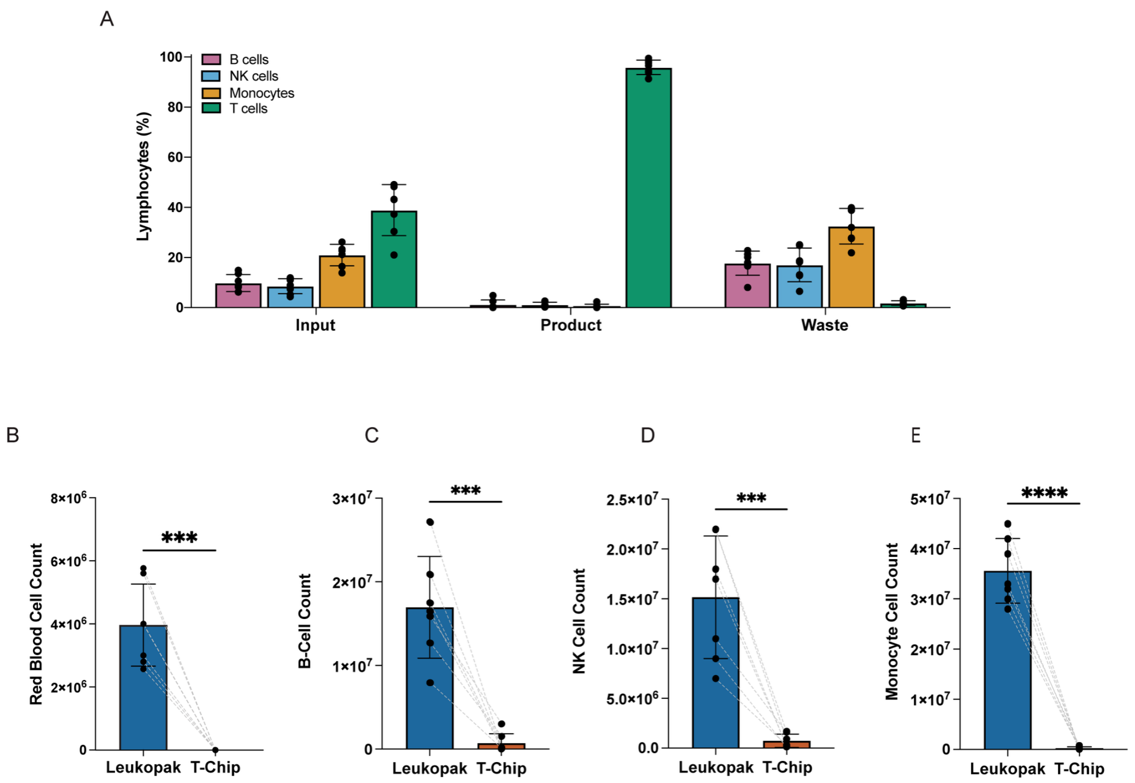


**Figure S2. Enrichment of T cells and removal of contaminating cell populations. (A)** Bar graphs comparing lymphocyte population distributions in the input leukopak mimic sample, the T-Chip product, and the T-Chip waste ($n$ = 7). T cells are enriched with minimal contamination. Error bars represent mean ± SD. T-Chip isolation depletes blood cell subpopulations, achieving **(B)** 5.0 ± 0.3 $Log_{10}$ RBC depletion ($n$ = 7), $p$ = 0.0002, **(C)** 2.0 ± 1.0 $Log_{10}$ B cell depletion ($n$ = 7), $p$ = 0.0005, **(D)**1.5 ± 0.4 $Log_{10}$ NK cell depletion ($n$ = 7), $p$ = 0.0007, **(E)** and 2.3 ± 0.5 $Log_{10}$ monocyte depletion ($n$ = 7), $p$ < 0.0001. Error bars represent the mean ± SD.

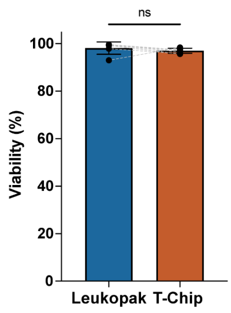


**Figure S3. Comparison of cell viability in the starting material leukopak and in the T-cell product after T-Chip enrichment.** T cells enriched using T-Chip maintain similar viability (97.0 ± 1.1%, $p$= 0.382, $n$ = 7), compared to that observed in an unprocessed leukopak mimic sample (98.1 ± 2.4%). Error bars represent mean ± SD.

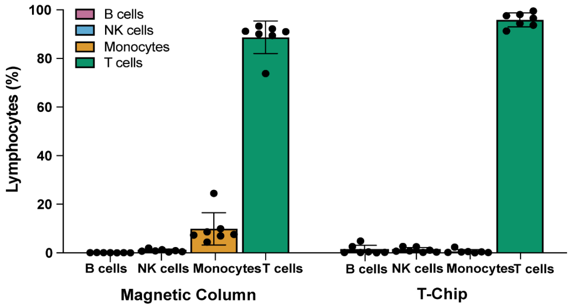


**Figure S4. Enrichment of T cells from leukopak mimic samples using Miltenyi magnetic column and T-Chip**. Bar graphs comparing lymphocyte population distributions in magnetic column and T-Chip products show that T-Chip enriches T cells with minimal contamination of B cells, NK cells, or monocytes (*n* = 7). Error bars represent mean ± SD.

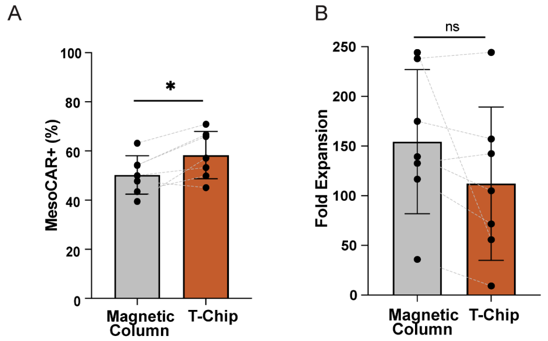


**Figure S5. Transduction efficiency of MesoCAR and T-cell expansion after isolation by magnetic column or T-Chip. (A)** MesoCAR transduction (*n* = 7) and **(B)** fold expansion of T cells isolated with a magnetic column and T-Chip on Day 13 (*n* = 7). Error bars represent mean ± SD.

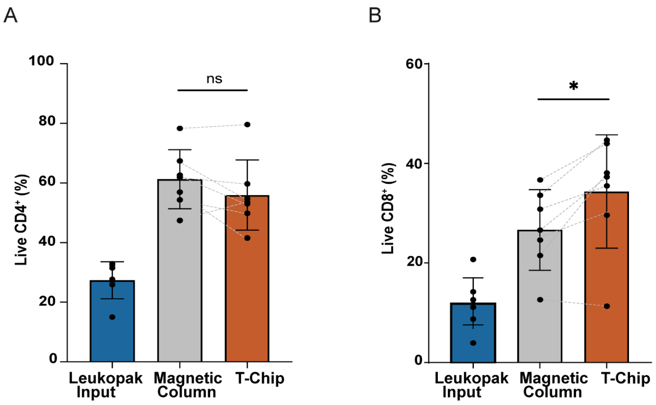


**Figure S6. Composition of CD4$^+$ and CD8$^+$ T cells after isolation using a magnetic column and T-Chip. (A)** Live CD4$^+$ and **(B)** CD8$^+$ T cells as percent of total cells in leukopak mimic, and in magnetic column and T-Chip products ($n$ = 7). Error bars represent mean ± SD.

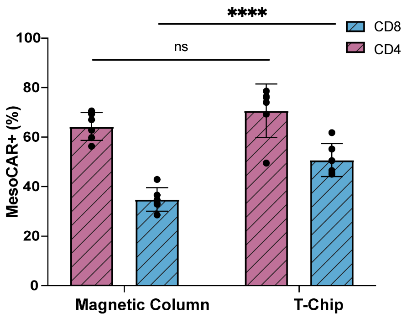


**Figure S7. MesoCAR transduction in CD4⁺ and CD8⁺ T cells.** Comparison of the transduction of cells isolated with a magnetic column and a T-Chip (*n* = 7). Error bars represent mean ± SD.

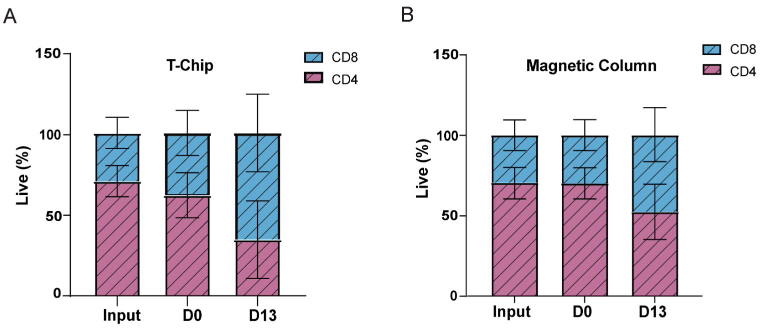


**Figure S8. Composition of T cells during the 13-day expansion.** Live $CD4^+$ and $CD8^+$ T cells in input leukopak, and in **(A)** T-Chip and **(B)** magnetic column products over a 13-day period ($n$ = 7). Error bars represent mean ± SD.

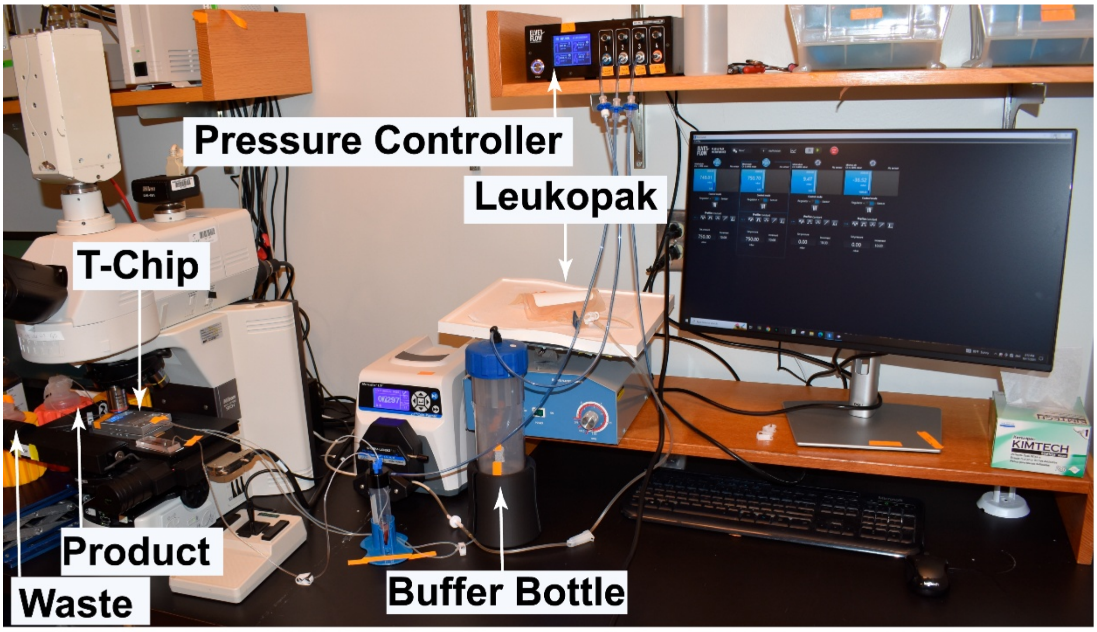


**Figure S9.** Experimental setup for clinical-scale T-cell isolation from large-volume leukapheresis products.

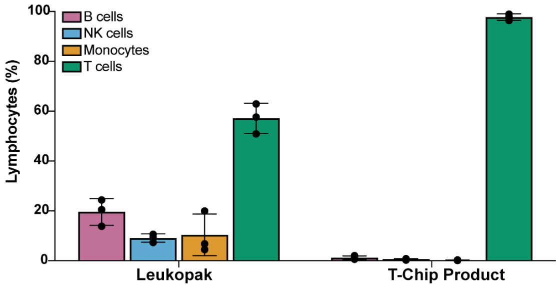


**Figure S10. T-Chip enrichment of T cells from large-volume leukopak samples.** (**A**) Bar graphs comparing lymphocyte population distributions in the input leukapheresis and T-Chip products ($n = 3$). (**B**) The T-Chip achieves excellent recovery of a highly pure T-cell population with high viability ($n = 3$). Error bars represent mean ± SD.

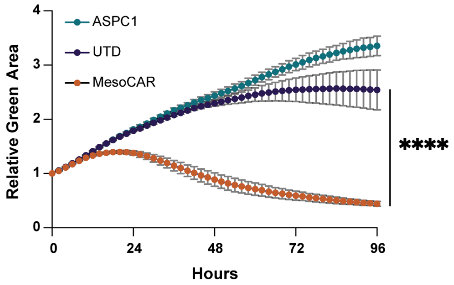


**Figure S11. Killing of GFP-ASPC1 pancreatic tumor cells over a 96-hour period by the MesoCAR T cells and donor-matched untransduced (UTD) T cells from leukopak samples compared to a tumor only control**. The relative green area represents tumor cell GFP fluorescence, measured by Incucyte live-cell imaging and normalized to t = 0 hours ($n$ = 3). Error bars represent mean ± SD.

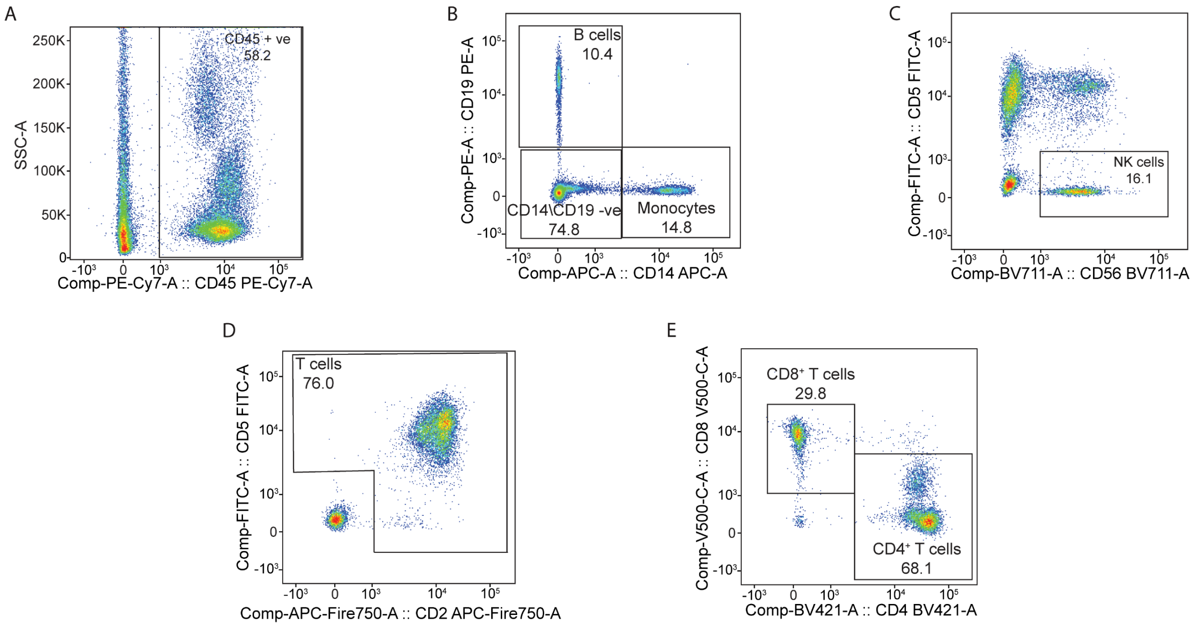


**Figure S12. Representative flow cytometry gating strategy of immune cell characterization panel in leukopak mimic input**. Phenotypic profiling was conducted to identify **(A)** immune cells, **(B)** B cells and monocytes, **(C)** NK cells, **(D)** T cells, and **(E)** T-cell subpopulations.

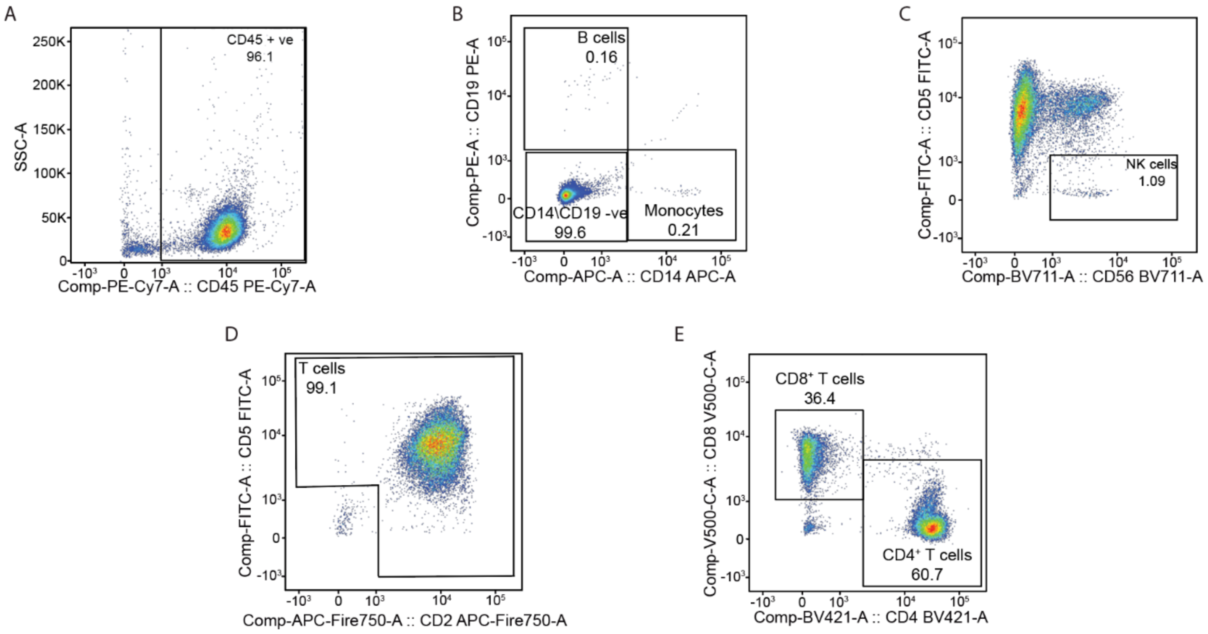


**Figure S13. Representative flow cytometry gating strategy of the immune cell characterization panel in the processed T-Chip product**. Phenotypic profiling was conducted to identify **(A)** immune cells, **(B)** B cells and monocytes, **(C)** NK cells, **(D)** T cells, and **(E)** T-cell subpopulations.

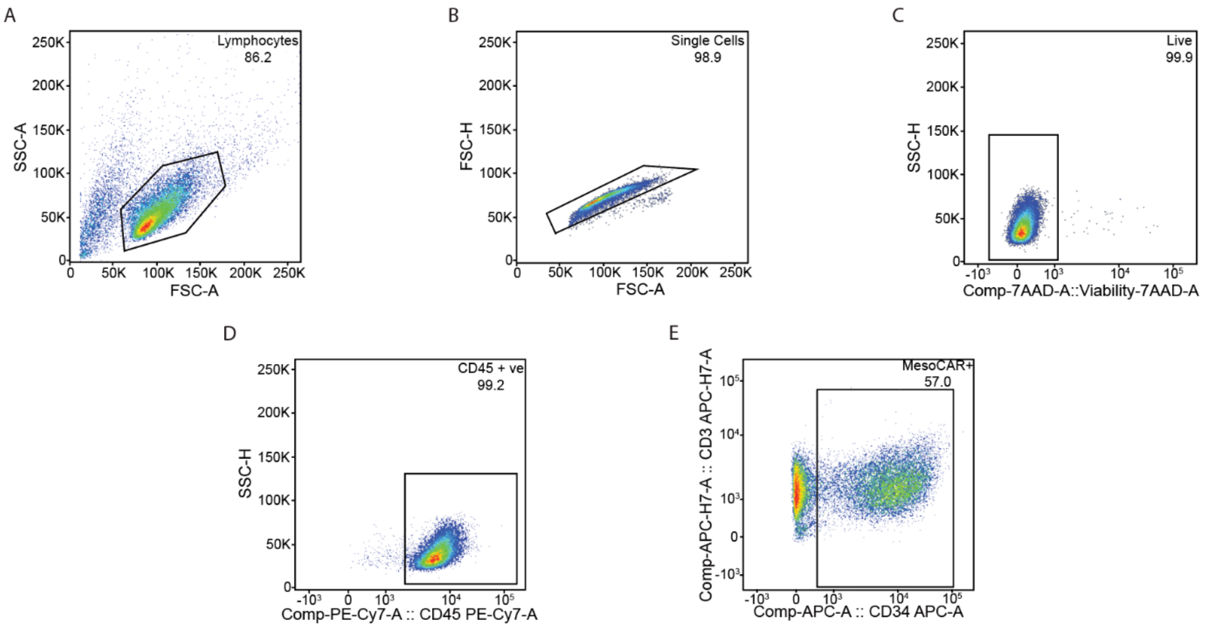


**Figure S14. Flow cytometry gating strategy of MesoCAR T cell characterization panel in CAR T cells manufactured from the T-Chip product**. Phenotypic profiling was conducted to identify **(A)** nucleated cells, **(B)** single cells, **(C)** live cells, **(D)** immune cells, and **(E)** MesoCAR+ cells.

**Supplementary Movie 1**

Sample video demonstrating the filling of T-Chip chambers with 40 µm soft iron particle suspension

**Supplementary Movie 2**

Computational simulation depicting the trajectory and magnetic deflection of an 8-µm T-cell labeled with a single magnetic bead within the T-Chip over a channel length of 20 mm.

**Supplementary Movie 3**

High-speed camera images show how $CD3^{+}$ T cells labeled with beads are selectively deflected into the central product channel in the T-chip's sorting channel, while unlabeled cells move into the waste channels.
